# Beta-Blocking Pavlov’s Bells: Propranolol Attenuates Compound Extinction in an Error-Dependent Manner

**DOI:** 10.64898/2025.11.29.691323

**Authors:** Ashleigh K. Brink, Clizia E. Martini, Laura H. Corbit

## Abstract

A significant limitation of extinction-based therapies is their failure to be expressed across time and different contexts. The re-emergence of the original behaviour provides evidence that extinction training does not erase the original learning but rather relies on new learning that suppresses the expression of the original behaviour. Thus, a strategy for reducing relapse phenomena is to enhance the inhibitory learning that occurs during extinction training so that extinction is more robust. One such strategy is compound extinction where a combination of previously reinforced stimuli are presented together for the first time during extinction training. This treatment has been shown to enhance extinction learning, evidenced by reduced future spontaneous recovery. However, the mechanisms are not fully understood. Experiment 1 assessed whether the compound extinction effect is the result of increased expectation of reward generated by the compound of stimuli that is then violated, driving further learning, or more simply, the novelty of the compound which reengages attention and thus promotes extinction without increasing prediction error. Experiment 2 tested whether the effects of the noradrenaline beta-receptor antagonist propranolol, shown elsewhere to reduce the compound extinction effect, relate to prediction error or novelty. We found that, when equating the novelty of the stimulus compound, larger prediction error resulted in better extinction evidenced as reduced spontaneous recovery. Further, we found that propranolol reduced this effect suggesting that prediction error rather than novelty is important for both the behavioural and pharmacological effects. Together our results point to behavioural and pharmacological strategies that can be used to improve the long-term expression of extinction.

**Highlights:**

- Increasing prediction error during extinction enhances extinction retention
- This effect is not explained by stimulus novelty
- Blocking beta-noradrenergic signaling attenuates the compound stimulus effect

## 1. Introduction

In Pavlov’s seminal 1927 experiments, he noted a phenomenon he would label *extinction*. He observed that presenting a conditional stimulus alone, in the absence of the reinforcer, would weaken performance of the conditional response. Pavlov also noted that evidence of the original learning persisted despite extinction, and could recover following the passage of time, a phenomenon he labelled *spontaneous recovery*. He would argue that during extinction the original learning was not permanently lost, a hypothesis which has garnered evidence over the past 100 years.

Spontaneous recovery (Rescorla, 1996, 1997, 2004), along with related phenomena like reinstatement (Rescorla & Heth, 1975) and renewal (Bouton & Bolles, 1979), have been incredibly influential in shaping the way people think about the learning that occurs in extinction (see Bouton et al., 2021; Quirk & Mueller, 2007). Together they suggest that extinction leaves intact at least a portion of the original learning and the suppression of performance seen across extinction training must occur through some process other than removal or erasure of that learning.

These demonstrations are important for understanding the mechanisms underlying extinction and its expression. However, in places where extinction is used as a means of reducing unwanted behaviour (e.g., exposure therapy) return of the original behaviour presents a significant obstacle to successful treatment. As such, there has been interest in identifying ways of augmenting or deepening extinction to improve its long-term expression(see Craske et al., 2014, 2022; Rosenberg et al., 2024).

One way of augmenting extinction learning is through compound extinction. In this paradigm, two previously rewarded stimuli are presented together followed by nonreinforcement, with the idea that the combination of stimuli should lead to greater expectation of reward. When that expectation is not met, this should produce greater learning than is achieved than if the stimuli are extinguished alone. Initially, Reberg (1972) noted that when two previously extinguished stimuli were paired, they could elicit an increase in responding compared to either stimulus alone. This increased responding provides evidence that animals’ expectations are based on the combination of stimuli with the predictive values summed or otherwise combined in some fashion. Next, Rescorla (2000) explored the longer term consequences of compound stimulus presentations. He observed that extinguishing two previously reinforced stimuli in compound enhanced retention of extinction learning compared to extinguishing a stimulus alone, or in compound with a neutral stimulus. Rescorla (2006) explored this compound extinction effect in depth, demonstrating similar summation during compound trials and deepened extinction of the constituent elements across multiple species (pigeons and rats), reinforcers (food reward or shock), and learning paradigms (Pavlovian and instrumental paradigms). In these experiments, Rescorla individually conditioned and extinguished three stimuli (A, X, Y) before presenting two of them together as a compound stimulus (AX) that underwent further extinction. He then went on to test responding to the remaining individual stimulus (Y) versus a stimulus extinguished in the compound (X) one week later. He found significantly strengthened extinction learning for the stimulus extinguished in compound (as demonstrated by reduced spontaneous recovery).

This approach has been replicated across a range of training conditions (Pavlovian, discriminated operant; spontaneous recovery, renewal) and types of reinforcers (food pellets, sucrose, cocaine, and alcohol; see Furlong et al., 2015; Janak et al., 2012; Janak & Corbit, 2010; Leung & Corbit, 2017).

While successful in reducing spontaneous recovery, the precise mechanism through which compound extinction is able to deepen extinction is not fully understood. While the idea that presentation of a compound of independently trained stimuli should serve to enhance prediction error and thus drive further learning has been offered as a likely explanation, alternative explanations for the effect exist. For example, the introduction of a stimulus compound during the course of extinction is novel and may itself be surprising to the animal. This could result in attentional changes and/or reengagement of learning without necessarily relying on generating a prediction error (or that prediction error might not be what drives learning).

While Rescorla (2000) offered some evidence against this, the scope of this effect has not been thoroughly explored. For example, if the benefit of compound extinction in reducing spontaneous recovery relies on prediction error, one might hypothesize that varying the magnitude of that prediction error should result in different degrees of spontaneous recovery, however, this has not been systematically investigated. The following experiments sought to examine these issues further. Experiment 1 trained rats with multiple discriminative stimuli that predicted either reward availability or its absence. During extinction, all groups were presented with compound stimuli, thus equating the novelty of encountering a stimulus compound. However, the degree to which those stimulus combinations predicted reward availability, and consequently, the degree to which that expectation was violated when responding was not reinforced, differed across groups. Therefore, this design allowed us to test whether the magnitude of the prediction error generated in compound extinction is important for improving extinction and reducing future spontaneous recovery. If prediction error is critical for augmenting extinction, greater error should result in greater extinction measured as reduced spontaneous recovery.

## 2. Materials & Methods

All procedures were conducted in accordance with ethical standards and guidelines established by the Canadian Council on Animal Care and protocols approved by the Animal Care Committee at the University of Toronto.

### 2.1. Experiment 1: The Benefit of Compound Extinction Effect is Related to Prediction Error

#### 2.1.1. Subjects

Subjects were 44, male Long-Evans rats (Charles River, St. Constant, QC, Canada). Animals were pair-housed in a temperature and humidity-controlled colony room on a 12-hour light-dark cycle (lights on at 7 AM). Experiments were conducted in the light phase. Rats were food restricted to maintain them at or above 90% of their free feeding weight throughout the experiment. Water and environmental enrichment were provided *ad libitum* in the home cage throughout the duration of the experiment.

#### 2.1.2. Apparatus

Training and testing took place in 16 identical behavioural chambers (Med Associates, Fairfax, VT, USA) housed individually within sound and light-attenuating cabinets. Each chamber contained a recessed magazine that included an infrared beam that when interrupted, recorded entries to the magazine, as well as two key lights situated above retractable levers and a magazine where 45-mg grain pellets could be delivered (F0165; BioServ, Flemington, NJ, USA). A white noise and 5-Hz clicker were used as auditory stimuli in addition to the visual key lights. The auditory stimuli were adjusted to 80 dB in the presence of background noise of 60 dB provided by a ventilation fan. Chambers were illuminated by a 3-W, 24-V house light. All experimental events were controlled and recorded with MED-PC V software (Med Associates).

#### 2.1.3. Magazine & Lever Training

Initially, rats received one 30-minute magazine training session where pellets were delivered on a random time 60 second schedule (RT-60s). On the following day, rats were trained to lever press for a food pellet reward on a continuous reinforcement schedule (one lever press earned one pellet). The session ended after 60 minutes or when 50 pellets were earned.

#### 2.1.4. Discrimination Training

Training going forward consisted of a discriminated operant procedure, in which rats were trained that responding during a 30-second stimulus was reinforced whereas responding in the absence of a stimulus was not. Each stimulus type (clicker, white noise, or light) was presented 8 times per session, for a total of 24 trials. During each auditory stimulus trial, lever pressing resulted in pellet delivery according to ratio schedules described below. Sessions also contained light stimulus trials, the meaning of which differed by group. Rats were randomly assigned to groups where the light functioned as either a discriminative stimulus (DS)+ (hereafter referred to as the DS+ condition; see Fig. 1A), a DS-condition where non-reinforced stimulus presentations were included in sessions throughout training, or a Habituation condition where the DS-was introduced on the last day of training to reduce its novelty. The limited training in this condition reduced the opportunity for an inhibitory association to form. Finally, a fourth group of animals (Inhibitor condition) received the light stimulus trained as a conditioned inhibitor. That is, the light stimulus was only presented in compound with one of the auditory stimuli, and its presence indicated responding would not be rewarded. Responding during that same auditory stimulus (noise or clicker, counterbalanced) was reinforced on trials where that stimulus was presented alone. The intertrial interval (ITI) varied between trials but averaged 90 seconds. Rats were trained on increasing random ratio (RR) schedules across days (Days 1-3: RR1; Days 4-6: RR2; Days 7-9: RR4).

**Fig. 1.**
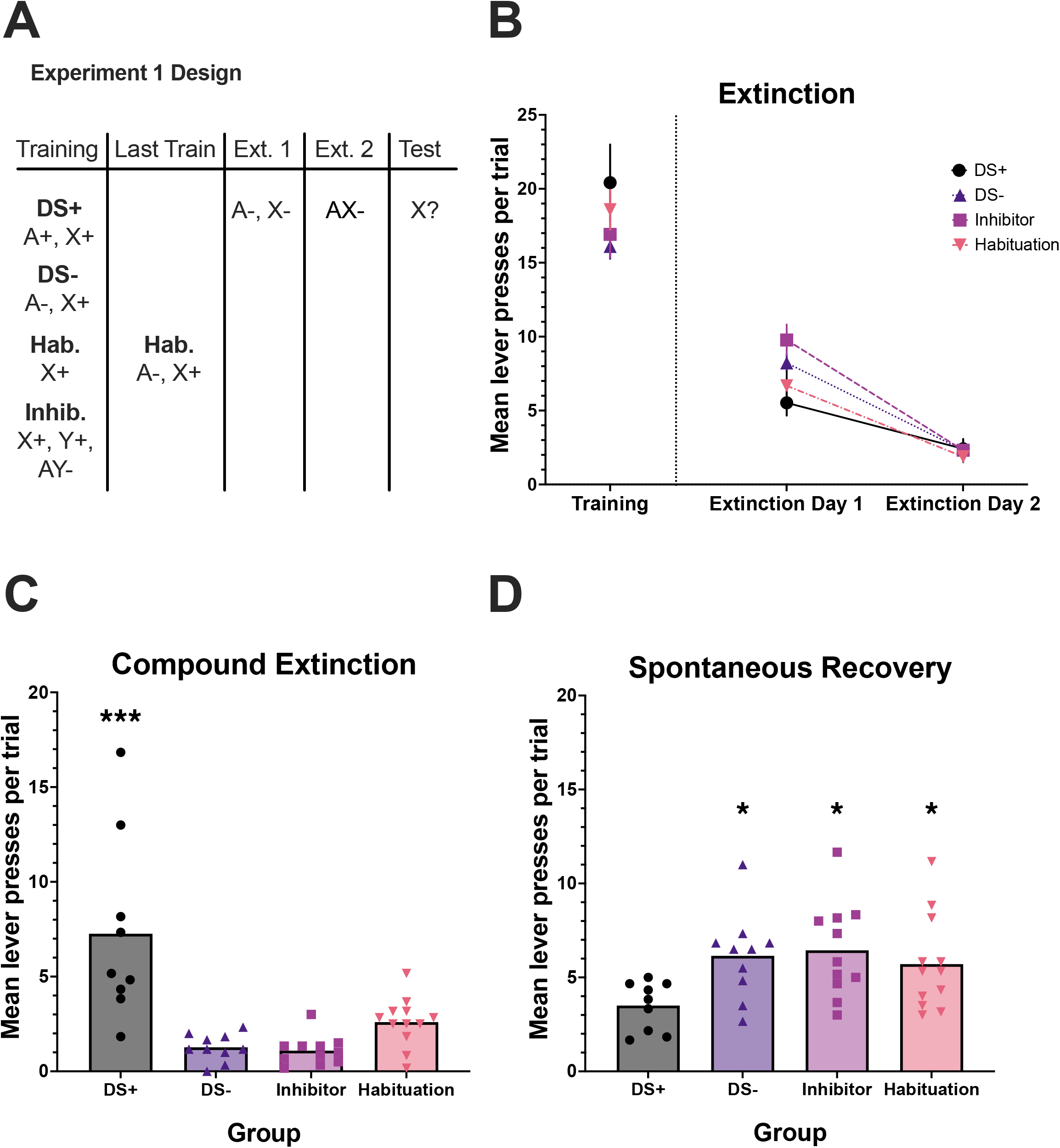
The compound extinction effect relies on the degree of prediction error associated with the compound. (**A**) Schematic representation of the study design. Rats were divided into one of four groups which received slightly different training. All rats underwent discriminated operant training where responding during an auditory cue, X, was reinforced. The consequence of responding during a visual cue, A, differed between these groups. Rats in the DS+ condition were reinforced during presentations of A. Rats in the DS-condition were not reinforced during presentations of A. Rats in the Habituation condition were also not reinforced after presentations of A, however, this stimulus was only introduced on the final day of training to decrease its novelty. All other groups maintained their previous training schedule on the final day. Rats in the Inhibitor condition received A trained as a conditioned inhibitor. That is, it was presented in compound with X, and on these trials, X was not reinforced. X alone trials were reinforced. No group differences were observed on the final training day (**B**, left panel; p’s > .05), or over the course of the first two extinction days (**B**, right panel; p’s > .05). On the compound extinction day all groups received AX trials that were not reinforced. Rats in the DS+ condition responded significantly more than rats in the other three conditions (**C**; p’s < .001). There was no difference in responding between the Habituation, Inhibitor and DS-conditions (p’s > .05). At test (**D**), the opposite pattern was observed. Responding was significantly lower in the DS+ group than the other 3 conditions (DS+ compared to Habituation, p = .033; Inhibitor, p = .006; DS-, p = .015). There was no difference in responding between the Habituation, Inhibitor and DS-conditions (p’s > .05). The results suggest that the compound extinction effect is reliant on the degree of error of the compound rather than the novelty of a compound stimulus. Error bars represent ± 1 SEM. Each dot in **C** and **D** represents one rat. ^*, **, ***^ = p < .05, p < .01, p < .001.

#### 2.1.5. Extinction Phase 1

On the next 2 days rats received training sessions that had 24 stimulus trials identical to training, but with the exception that no pellets were delivered.

#### 2.1.6. Extinction Phase 2: Compound Extinction

Rats received 6 presentations of a compound stimulus (Light + Noise or Clicker, counterbalanced) under extinction. For the Inhibitor group, animals received the auditory stimulus that had not been paired with the light in compound trials in training.

#### 2.1.7. Spontaneous Recovery Test

Four weeks later, rats were assessed for retention of extinction. The auditory cue extinguished in compound was presented 6 times. No pellets were delivered in this session. We hypothesized that we would see the best retention of extinction in this test in the DS+ condition compared to the other three conditions, as it encountered the largest prediction error corresponding to presentations of the compound stimulus.

### 2.2. Experiment 2: Propranolol Reduces the Benefit of Extinction of a Compound of Two Excitatory Stimuli

There are now multiple demonstrations that blocking noradrenergic beta receptors during compound extinction limits the ability of this manipulation to enhance extinction (Furlong et al., 2015; Janak et al., 2012; Janak & Corbit, 2010; Leung & Corbit, 2017). While Experiment 1 confirmed that the magnitude of the prediction error generated during compound extinction contributes to the strength of subsequent extinction learning, the exact nature of the interaction between noradrenaline and error in this paradigm remained unexplored. Considering that noradrenergic locus coeruleus neurons respond to both novelty (Fois et al., 2022; Sara et al., 1994; Sara & Segal, 1991; Takeuchi et al., 2016; Vankov et al., 1995) and the omission of expected rewards (i.e., negative prediction error; Sara et al., 1994; Sara & Segal, 1991; Su & Cohen, 2022), Experiment 2 sought to test whether the impact of beta-receptor antagonism differed according to the degree of prediction error generated by the stimulus compound. If the large prediction error generated when a compound of two previously reinforced stimuli is not reinforced recruits noradrenaline release, treatment with the beta-receptor antagonist propranolol should weaken the compound extinction effect (i.e., allow greater spontaneous recovery) despite the large prediction error. In contrast, minimal recruitment of noradrenaline and thus limited effects of beta-receptor antagonism should be expected when prediction error is small. Conversely, if it’s the novelty of the compound that recruits noradrenaline release and drives learning, this should be similar for any novel compound and independent of the magnitude of the prediction error. If so, propranolol should weaken extinction relative to saline-treated animals and there should be no difference between groups where the error generated by the compound is large versus small.

#### 2.2.1 Subjects

Forty (20 male, 20 female) Long-Evans rats served as subjects (Charles River). They were trained and tested in two cohorts (n = 24 Cohort 1, n = 16 Cohort 2). Animals were housed and maintained in the same conditions described in Experiment 1.

#### 2.2.2. Apparatus

Training and testing took place in operant chambers identical to those described in Experiment 1.

#### 2.2.3. Magazine & Lever Training

This phase of training was identical to Experiment 1.

#### 2.2.4. Discrimination Training

Further training consisted of a discriminated operant procedure, in which rats were trained to respond during a 30 second presentation of an auditory or visual stimulus (24 trials total, 8 of each clicker, white noise, and light). During each auditory stimulus, lever pressing resulted in pellet delivery. Rats were randomly assigned to one of two groups where the light served as either a DS+ or DS-(DS+, DS-conditions from Experiment 1; see Fig. 2A). The ITI averaged 90 seconds. Rats were trained on increasing random ratio (RR) schedules across groups of 3 days (Day 1-3: RR1; Day 4-6: RR2; Day 7-9: RR4). Rats received one mock injection of saline during the final day of training.

**Fig. 2.**
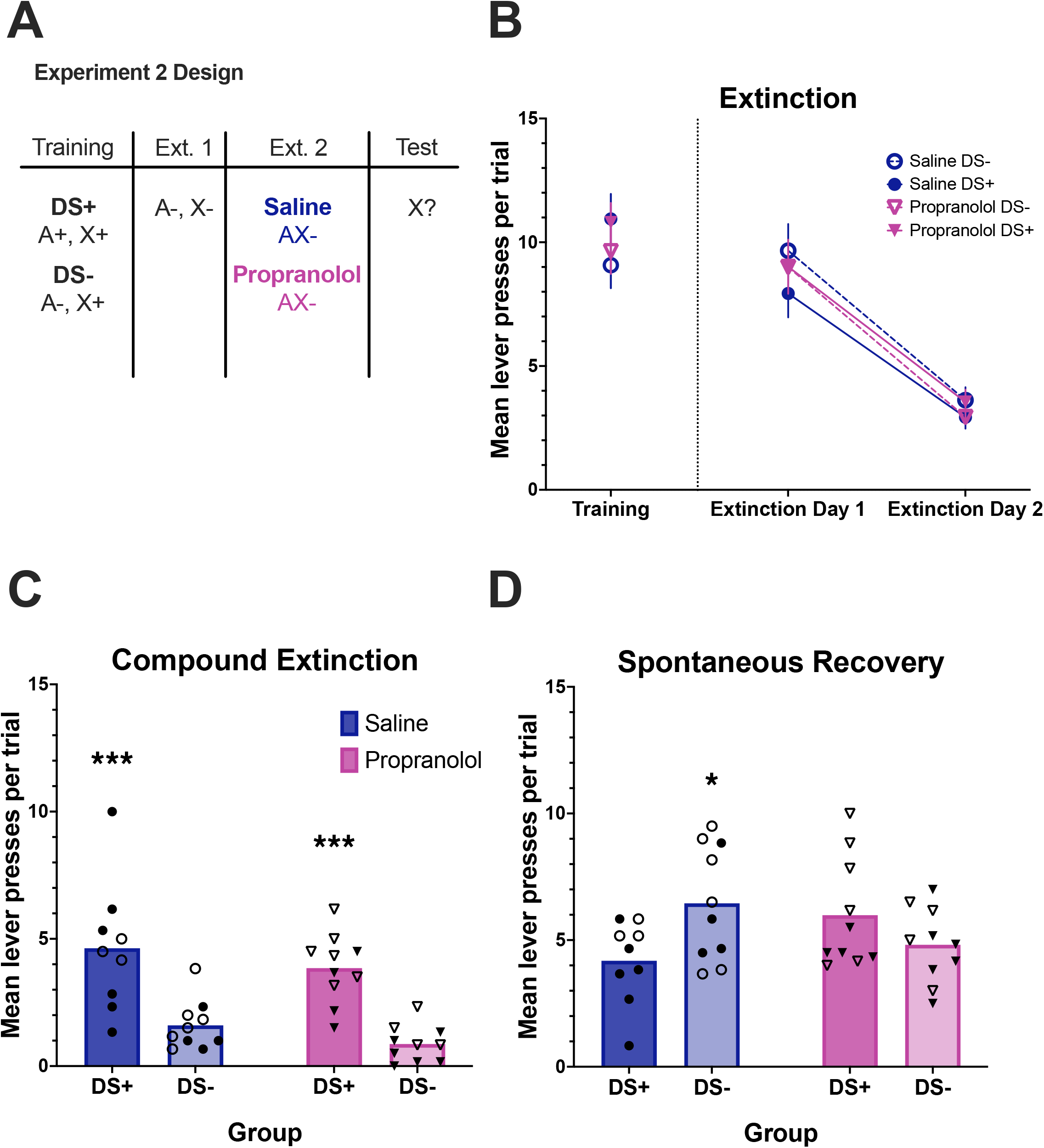
Noradrenergic signalling is recruited in response to high-error but not low error compound extinction. (**A**) Schematic representation of the study design. Rats were randomly assigned into either the DS+ or DS-group (trained as in Experiment 1). Prior to the compound extinction day these rats were further counterbalanced into Propranolol and Saline conditions, giving a total of four, between-subject groups. There were no differences in responding between groups on the final day of training (**B**, left panel; p’s > .05) or on the first two days of extinction (**B**, right panel; p’s > .05). Thirty minutes prior to the compound extinction session, rats received an IP injection of either sterile saline or propranolol (10 mg/kg). During the compound session, rats in the DS+ conditions responded significantly more than rats in the DS-conditions (**C**; p < .001). There was no effect of drug condition (p = .151). At test (**D**), there was no significant effect of drug condition or compound type (p’s > .05). Notably, the female rats responded more than the male rats across the entire test (p = .040), however this did not interact with any other factor (p’s > .05). Importantly, there was an interaction between drug and compound type (p = .033), with follow-up analyses indicating a significant effect of compound type across the saline animals (p = .045), but not the propranolol animals (p = .306). This suggests that noradrenaline is necessary for the compound extinction effect under high error but not low error conditions. Error bars represent ± 1 SEM. Each dot in **C** and **D** represents one rat. Filled dots represent male rats, empty dots represent female rats. ^*, **, ***^ = p < .05, p < .01, p < .001.

#### 2.2.5. Extinction Phase 1

All rats received two extinction sessions as in Experiment 1.

#### 2.2.6. Extinction Phase 2: Compound Extinction & Drug Administration

Prior to this session, rats from each group were further divided into Saline and Propranolol groups. Rats received 6 presentations of a compound stimulus (Light + Noise or Clicker, counterbalanced within group) under extinction. Thirty minutes prior to this final extinction session, rats received an intraperitoneal injection of either saline or propranolol hydrochloride (10 mg/kg; Sigma-Aldrich, Oakville, ON, Canada). We hypothesized that propranolol would attenuate the effect of compound extinction in the DS+ condition but leave compound extinction in the DS-conditioned unaffected.

#### 2.2.7. Retention Test

On the day following compound stimulus presentation and drug administration, rats were assessed for their learning. The auditory cue extinguished in compound was presented 3 times. No pellets were delivered in this session.

#### 2.2.8. Spontaneous Recovery Test

Four weeks later, animals were assessed for the return of responding over time. The session was identical to the probe test, except it was 6 trials.

## 3. Results

### 3.1. Experiment 1: The Benefit of Compound Extinction Effect is Related to Prediction Error

The aim of this experiment was to assess whether the magnitude of the prediction error generated by different combinations of stimuli, rather than the novel introduction of a compound well into extinction training, was responsible for the benefit to extinction previously observed with compound extinction.

#### 3.1.1 Experiment 1: Training

On the final day of acquisition there were no group differences in responding to the auditory stimulus that would ultimately be tested for spontaneous recovery following compound extinction [target stimulus; F(3, 38) = 0.743, p = .533].

#### 3.1.2 Experiment 1: Initial Extinction

Responding decreased over the initial two days of extinction [F(1, 38) = 140.1, p = .001]. Prior to the differential treatment, there were no group differences in responding to the target stimulus [F(3, 38) = 1.184, p = .329]. There was no interaction between group and extinction day [F(3, 38) = 2.225, p = .101].

#### 3.1.3 Experiment 1: Compound Extinction

On the third day of extinction, compound trials were introduced to each group. As shown in Fig. 1C, rats that received a compound comprised of two excitors (DS+ condition) responded robustly in these trials. Groups that received a compound including either a pre-exposed stimulus (i.e., Habituation, DS-) or one that was trained as a conditioned inhibitor (Inhibitor condition) responded much less to the compound, despite it being novel in all conditions. Importantly, this pattern reversed in the test of the target stimulus conducted four weeks later; responding to the stimulus extinguished in compound with a second excitor was less than in any of the other conditions. This description is supported by the statistical analyses which revealed a significant effect of experimental phase [extinction vs. test; F(1, 38) = 22.588, p = .001], no effect of group [F(3, 38) = 2.173, p = .107], but a significant interaction between phase and group [F(3, 38) = 15.760, p = .001].

#### 3.1.4 Experiment 1: Test

Because the rats received a stimulus compound in the extinction phase and were tested with a single stimulus in the spontaneous recovery test the amount of responding across phases is expected to differ; therefore, in order to further evaluate the effect of the magnitude of prediction error in producing the observed effects, we focused on between-group rather than within group comparisons. Simple effects analyses demonstrated that for the extinction day, the identity of the compound determined how much responding was observed [F(3, 38) = 14.064, p = .001]. Post-hoc analyses (LSD) indicated that rats in the DS+ condition responded more than those in any other condition [all p’s < .001] which did not differ from each other [all p’s > .05]. During the spontaneous recovery test there was again a significant effect of group [F(3, 38) = 3.274, p = .031] and post-hoc analyses demonstrated that responding in the DS+ group was significantly lower than in the Habituation group [p = .033], the Inhibitor group (p = .006) or the DS-group (p = .015). Responding between these latter three groups did not differ [all p’s > .05]. Together these results demonstrate that the benefit to the long-term expression of extinction found following compound extinction relies on the degree to which that compound produces a prediction error and that despite the novelty of other compounds, introduction of additional stimuli that do not also increase reward expectancy has little effect on extinction.

#### 3.2.1 Experiment 2: Propranolol Reduces the Benefit of Extinction of a Compound of Two Excitors

The aim of this experiment was to assess what recruits noradrenergic signalling in the compound extinction paradigm – the magnitude of the prediction error generated by the compound stimulus, or the novel introduction of a compound well into extinction training,

#### 3.2.2 Experiment 2: Training

To ensure equal responding between groups at the end of training (Fig. 2B), a one-way ANOVA was conducted. There was no effect of future drug condition [F(1,31) = 0.003, p = .959], compound type [F(1,31) = 3.526, p = .070], sex [F(1,31) = .078, p = .782], or sex x drug interaction [F(1,31) = 1.326, p = .258]. No other interactions between these factors were significant [F’s < 1].

#### 3.2.3 Experiment 2: Initial Extinction

The initial two extinction days were assessed with a mixed model ANOVA. Responding significantly decreased across days [F(1, 31) = 168.915, p < .001]. There was no significant effect of sex [F(1,31) = 2.803, p = .104], compound type [F(1,31) = 0.357, p = .554], future drug condition [F(1,31) = 0.025, p = .876], or day x sex interaction [F(1,31) = 1.968, p = .171]. No other interactions between these factors were significant [F’s < 1].

#### 3.2.4 Experiment 2: Compound Extinction

During the compound extinction day (Fig. 2C), there was no significant difference in responding due to drug condition [F(1,31) = 2.173, p = .151] or sex [F(1,31) = 1.900, p = .178]. However, there was a significant effect of compound type [F(1,31) = 32.706, p < .001], with the DS+ groups responding significantly more than the DS-groups. This indicates successful summation – the animals in DS+ groups had higher expectation of reward than animals in DS-groups. No interactions within these variables were significant [F’s < 1].

#### 3.2.5 Experiment 2: Test

In the test of spontaneous recovery conducted 4 weeks later (Fig. 2D), there was no significant difference in responding by compound type [F(1,31) = 0.626, p = .435] or drug condition [F(1,31) = 0.028, p = .868]. There was a significant difference in responding by sex [F(1,31) = 4.578, p = .040], which represents broadly heightened responding in females versus males. Importantly, as hypothesized, there was a significant compound type x drug condition interaction [F(1,31) = 4.967, p = .033]. Simple main effects analyses used to interpret this interaction indicated a significant effect of compound type in saline animals [F(1, 31) = 4.353, p = .045], but not in propranolol animals [F(1, 31) = 1.085, p = .306]. There were no interactions between compound type and sex [F(1,31) = 1.028, p = .318], drug and sex [F(1,31) = 0.001, p = .974], or compound type, drug, and sex [F(1,31) = 0.050, p = .825].

The saline groups’ results are consistent with what was observed in drug-free animals in Experiment 1 – spontaneous recovery is significantly higher in the DS-group than the DS+ group. This again confirms that the compound extinction effect is driven by error, rather than the novelty or salience of the compound stimulus itself. The propranolol groups however show no difference in spontaneous recovery. This suggests that the attenuation of noradrenergic signalling attenuated DS+ compound extinction as previously shown (e.g., Janak & Corbit, 2011). Importantly however, there was no effect on extinction of a DS-compound. Ultimately, this indicates that noradrenergic activity is recruited under high error conditions (DS+), but not low error (DS-) conditions.

## 4. Discussion

In these experiments we provide further insight into the mechanisms underlying the compound extinction effect and the nature of the contribution of noradrenaline to promoting lasting extinction. Experiment 1 shows that augmenting prediction error during extinction training promotes lasting extinction. Extinguishing a compound of two previously reinforced stimuli (DS+ condition), but not compounds containing inhibitory (Inhibitor condition), ostensibly neutral (DS-condition), or habituated (Habituation condition) stimuli resulted in less responding to the element of the compound (X) tested alone four weeks later. This suggests that the ability of a stimulus compound to improve the long-term expression of extinction depends on the degree to which the stimuli that comprise that compound led to an expectation of a reinforcer that is then omitted.

The increase in responding seen to the compound of previously reinforced stimuli suggests that animals sum the values expected based on each stimulus present during compound trials (Pavlov, 1927; Reberg, 1972; Rescorla, 2006; Weiss, 1964). This can account for why responding to the DS+ compound was greater than to the other compounds. The enhanced error term can then drive further learning (Rescorla, 2006). However, this raises the question of whether it’s the violation in expectations based on the presence of stimuli that drives learning, or whether the increase in responding to the compound which, in the discriminated operant paradigm, provides additional opportunities to learn that responding is not reinforced thus leading to greater extinction.

Although not directly tested here, several previous findings suggest that it is the former. Rescorla (2006) took advantage of the fact that pigeons respond differently to localized versus diffuse stimuli. When trained with a localized stimulus such as a key light paired with reward, pigeons will peck as a conditional response. If a diffuse stimulus such as a white noise is paired directly with grain reward this produces excitation but does not increase pecking. In contrast, if a diffuse stimulus (A) is trained as a facilitator (X-, AX+) this generates a stimulus that enhances responding (i.e., A enhances responding to X) but is not itself excitatory. To dissociate the effects of excitation and enhanced responding Rescorla (2006; Experiment 5) trained pigeons with localized visual stimuli that predicted reward. The animals were also trained with two diffuse stimuli, one of which was trained as an excitor and the other trained as a facilitator. The localized stimuli then underwent extinction alone followed by a phase where compound trials were introduced where one compound contained the diffuse excitor and one contained the facilitator. During these compound trials, as anticipated, the diffuse facilitator in compound with the localized stimulus increased responding whereas the compound containing the diffuse excitor did not. Importantly, when the target stimuli were tested one week later there was less responding to the stimulus that had been extinguished alongside the excitor. Thus, these results suggest that the presence of excitation rather than increased responding is critical for enhanced extinction.

Related findings come from experiments where atomoxetine, a noradrenaline reuptake inhibitor (Janak et al., 2012; Janak & Corbit, 2010; Leung & Corbit, 2017) or optogenetic stimulation of the locus coeruleus (Lui et al., 2024), the primary source of noradrenaline to the forebrain (Poe et al., 2020), were administered during extinction of a single stimulus. While neither treatment increased responding relative to control conditions, each enhanced the long-term retention of extinction. While these results don’t rule out the possibility that more responding allows more opportunity for learning, they do demonstrate that an increase in responding is not required to augment behavioural extinction.

In Experiment 2 we again examined the role of noradrenaline in compound extinction. The observed results suggest that noradrenaline contributes to the enhanced extinction resulting from compound extinction under high error conditions (DS+), but not low error (DS-) conditions. This builds upon previous work (Furlong et al., 2015; Janak et al., 2012; Janak & Corbit, 2010; Leung & Corbit, 2017), which has indicated that systemic manipulations of noradrenaline can bidirectionally affect extinction, with heightened noradrenaline activity strengthening extinction of a single stimulus, and attenuated noradrenaline activity impairing compound extinction. As mentioned above, these studies also examined whether propranolol would attenuate extinction of a single cue. They found that propranolol had no effect, mirroring our low error compound (DS-) results. However, unlike the current experiments, these previous studies did not control for the novelty of the compound stimulus itself.

More recent work has focused on the epicentre of noradrenergic activity in the brain, the locus coeruleus. As with the above systemic investigations into noradrenaline and extinction, it has been demonstrated that optogenetic stimulation of the locus coeruleus enhances extinction of a single cue, whereas inhibition had no effect (Lui et al., 2024). In another recent study, it was found that pharmacological inhibition of the locus coeruleus attenuated compound extinction, as well as another related phenomenon known as overexpectation, in which reward is less than expected (Brink et al., 2025). Notably, Brink et al. (2025) also examined high vs. low error compounds, in a within-subjects design and found concordant results. In combination with the presently reported findings, a clear picture is beginning to emerge in our understanding of the compound extinction paradigm, and further, the role of noradrenaline in this paradigm.

In summation, this study provides novel evidence that the compound extinction effect is driven by prediction error (i.e., a heightened prediction of reward), rather than the novelty of the compound stimulus. Further, we demonstrate that noradrenergic signalling is predominantly recruited in response to high error, but not low error compounds. These results provide a further foundation for work examining the role of noradrenaline in extinction, and more broadly, in error-driven learning.

## Abbreviations

(DS): Discriminative Stimulus

